# Extreme phylogenomic diversity in non-pathogenic *Xanthomonas* strains associated with citrus

**DOI:** 10.1101/835645

**Authors:** Kanika Bansal, Sanjeet Kumar, Prabhu B. Patil

## Abstract

*Xanthomonas* are primarily known as a group of phytopathogens infecting diverse plants. Recent molecular studies reveal existence of novel species and strains of *Xanthomonas* that follow non-pathogenic (NPX) lifestyle. Presently, we report whole genome sequence of four NPX reported from *Citrus* sp. (NPXc). Taxonogenomic analysis revealed surprising diversity as each of the three isolates were found to be novel genomospecies that together form a *Citri* associated non-pathogenic *Xanthomonas* species complex (NPXc complex). Interestingly, this NPXc complex is related to another non-pathogenic species, *X. sontii* from rice (NPXr). On the other hand, the fourth NPXc isolate was found to be related to NPX isolates from walnut (NPXw), altogether they form a potential taxonomic outlier of pathogenic *X. arboricola* species. Interestingly, their pathogenic counterparts, *X. citri* and *X. arboricola* are known to cause similar symptoms in diverse fruit plants. Further, genomic investigation of well-characterized pathogenicity clusters in NPXc isolates revealed lifestyle specific gene content dynamics. Majorly, genes essential for virulence (T3SS, effectors, T6SS, biofilm cluster etc.) and adaptation (*gum, rpf*, iron uptake and utilization, T2SS (*xps*) and variable repertoire of cell wall degrading enzymes) could be depicted by comparative genomics of *Xanthomonas* community comprising of diverse lifestyle isolates. Overall, present analysis confers that non-pathogenic isolates of diverse host phylogenomically converge and are evolving in parallel to their pathogenic counterparts. Hence, there is a need to understand the world of NPX isolates from diverse and economically important hosts. Genomic knowledge and resource will be invaluable in both basic and applied research of genus *Xanthomonas*.

**Impact statement:** *X. citri* is one of the top phytopathogenic bacteria and is the causal agent of citrus canker. Interestingly *Xanthomonas* is also reported to be associated with healthy citrus plants. Advent of genomic era is allowing us carry out detailed evolutionary study of *Xanthomonas* community associated with citrus and other plants. Our investigations have revealed hidden and extreme inter-stain diversity of non-pathogenic *Xanthomonas* strains from citrus plants warranting further large scale studies. This indicated unexplored world of *Xanthomonas* from healthy citrus plants species that may be co-evolving as a species complex with host unlike the variant pathogenic species. The knowledge and genomic resource will be valuable in evolutionary studies exploring its hidden potential and management of pathogenic species.

## Introduction

*Xanthomonas* is a complex phytopathogen with an array of host range including 268 dicots and 124 monocots (Leyns, De Cleene et al. 1984, Hayward 1993, Chan and Goodwin 1999). The genus comprises of 34 different species with diversifying pathogenic potential. *Citrus* sp. is widely known to be infected with *X. ctiri*, a quarantine pathogen (Gottwald, Hughes et al. 2001, Gottwald, Sun et al. 2002, Brunings and Gabriel 2003, Graham, Gottwald et al. 2004). Owing to its economic importance as one of the top phytopathogenic bacterium, its evolution into successful species is of interest to researchers, policy makers etc. Moreover, it is most successful *Xanthomonas* species harboring twenty-two pathovars known to be host specific to diverse plants (Bansal, Midha et al. 2017).

Interestingly, out of these 34 species, only two species i.e. *X. sontii* and *X. maliensis* are exclusively having non-pathogenic *Xanthomonas* strains associated with rice (NPXr), while one species i.e. *X. arboricola* is having strains with both pathogenic and non-pathogenic lifestyles on the walnut (Cesbron, Briand et al. 2015, Essakhi, Cesbron et al. 2015, Triplett, Verdier et al. 2015, Bansal, Kaur et al. 2019). In addition to these, there are various unclassified and unexplored non-pathogenic isolates reported from a diversity of host plants (Vauterin, Yang et al. 1996). These host plants are already associated with successful pathogenic species of *Xanthomonas* and reported to cause devastating disease worldwide. Though, non-pathogenic strains have significant impact on evolution of *Xanthomonas* genus as a whole, yet, they are widely overlooked due to their less economic importance.

Genomic era has radically transformed our basic understanding of bacterial evolution. Phylogenomics and taxonogenomics are revealing the evolutionary processes that lead to genome diversification and speciation of the order *Lysobacterales* comprising the genus *Xanthomonas* (Kumar, Bansal et al. 2019). *Xanthomonas* community analysis by comparative genomics of diverse lifestyles bacteria is crucial for our understanding. Lifestyle and adaptability of a microbe to a new host or environment depends upon its genomic content and potential. In the *Xanthomonas* genus, there are widely known virulence-related gene(s) or gene cluster(s) such as type secretion systems and their effectors (Jha, Rajeshwari et al. 2007, Lu, Patil et al. 2008, Büttner and Bonas 2010, Bernal, Llamas et al. 2018, Bansal, Midha et al. 2019), *rpf* gene cluster (Dow, Feng et al. 2000, Chatterjee and Sonti 2002), iron uptake, utilization and siderophore related genes (Etchegaray, Silva-Stenico et al. 2004, Pandey and Sonti 2010, Subramoni, Pandey et al. 2012), global regulators and two component systems (Wengelnik and Bonas 1996, Wengelnik, Rossier et al. 1999, Tsuge, Nakayama et al. 2006, Lee, Jeong et al. 2008, Qian, Han et al. 2008) etc. Status of these virulence related gene content amongst non-pathogenic strains of *Xanthomonas* will be of great significance. Genome-wide comparison of pathogenic and non-pathogenic isolates of *X. arboricola* reflects various genomic determinants like: T3SS, chemoreceptors, adhesins etc (Cesbron, Briand et al. 2015). Similarly, such an analysis of rice *Xanthomonas* community from our group have demonstrated gene(s) or gene cluster(s) essential for virulence and adaptation to the host (Bansal, Midha et al. 2019). Similarly, such a study on non-pathogenic *Xanthomonas* from diverse hosts like economically important citrus plants is lacking and is the need of the hour.

In the present analysis, we have focused on evolution of *Citrus* sp. associated *Xanthomonas* community, already known to have a global devastating pathogen. Along with that, we have also included genomes of NPX isolates from walnut and rice available in NCBI. Here, we have sequenced four NPXc strains and carried out phylogenomics, taxonogenomics and comparative genomics study of the NPX isolates. Primarily, we tried to address whether all these non-pathogenic counterparts from diverse hosts belong to a species, irrespective of their host of isolation or they belong to diverse species? What is their relation with two of the already known non-pathogenic species amongst *Xanthomonas* genus? What is genomic content and flux of gene(s)/gene cluster(s) during the course of evolution. Present study will be a genomic investigation towards understanding of widely unexplored non-pathogenic isolates and evolution of successfully established pathogenic *Xanthomonas* species.

## Results

### Whole genome sequencing and assembly of NPXc isolates

The genome of four NPXc were sequenced using Illumina MiSeq platform. The assemblies obtained were having coverage ranging from 73X to 146X with N50 values of 94 Kb to 213 Kb. Genome size and GC content of all the strains were approximately 5Mb and 68% respectively and genome statistics are summarized in table 1.

**Table 1:**
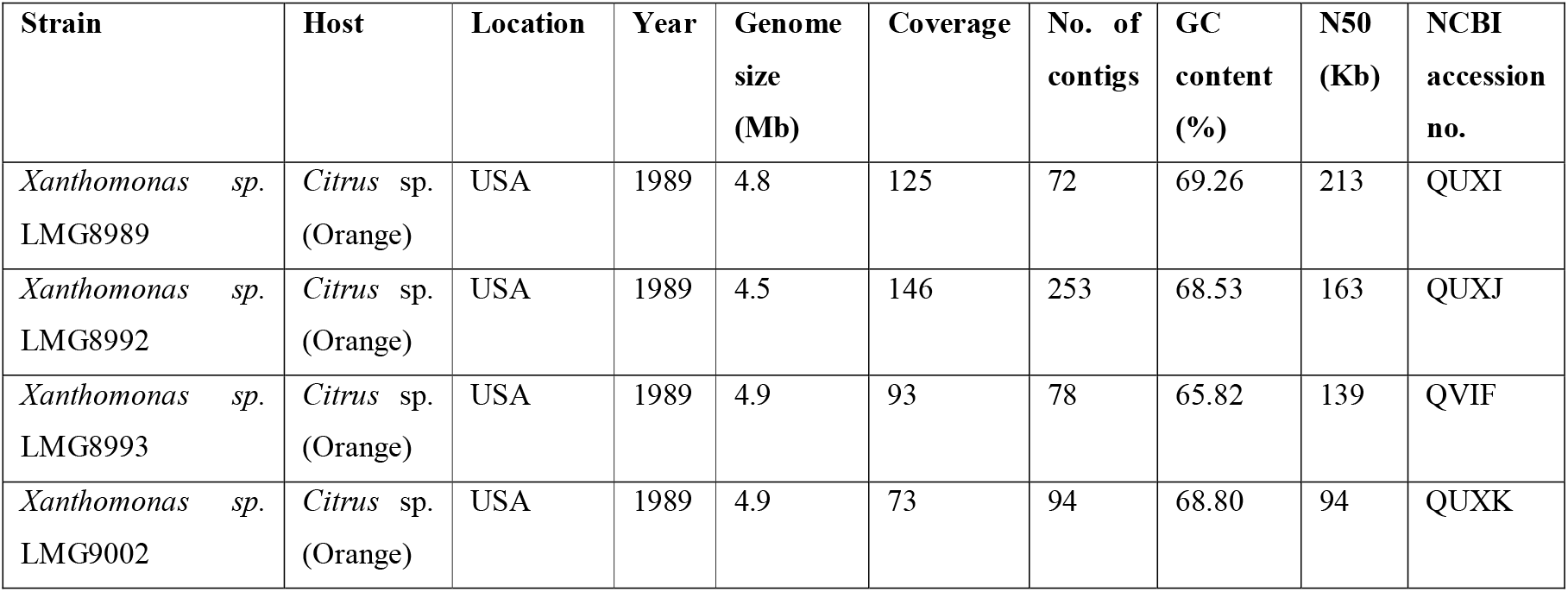
Assembly statastics and metadata of NPXc strains in the present study.

| Strain | Host | Location | Year | Genome size (Mb) | Coverage | No. of contigs | GC content (%) | N50 (Kb) | NCBI accession no. |
| --- | --- | --- | --- | --- | --- | --- | --- | --- | --- |
| *Xanthomonas* sp. LMG8989 | *Citrus* sp. (Orange) | USA | 1989 | 4.8 | 125 | 72 | 69.26 | 213 | QUXI |
| *Xanthomonas* sp. LMG8992 | *Citrus* sp. (Orange) | USA | 1989 | 4.5 | 146 | 253 | 68.53 | 163 | QUXJ |
| *Xanthomonas* sp. LMG8993 | *Citrus* sp. (Orange) | USA | 1989 | 4.9 | 93 | 78 | 65.82 | 139 | QVIF |
| *Xanthomonas* sp. LMG9002 | *Citrus* sp. (Orange) | USA | 1989 | 4.9 | 73 | 94 | 68.80 | 94 | QUXK |

### Phylogenomic and taxonogenomic diversity of NPX isolates

Phylogenomic analysis of available non-pathogenic isolates (supplementary table 1) along with their pathogenic counterparts and other type or representative strains is shown in figure 1. Majority of the NPX such as NPXc (LMG8989, LMG8992 and LMG9002) and NPXr (*X. sontii* PPL1, *X. sontii* PPL2, *X. sontii* PPL3, LMG12459, LMG12460, LMG12461, LMG12462, SHU166, SHU199, SHU308) belong to clade II of *Xanthomonas* genus. Whereas, NPXw (*X. arboricola* CFBP7634 and *X. arboricola* CFBP7651) and one of the NPXc (LMG8993) belong to clade I with *X. arboricola* as their closest phylogenomic relative. Further, *X. maliensis* (NPXr) was in between clade I and clade II (figure 1). Further, to confirm the species status of these isolates, taxonogenomic analysis, including ANI and dDDH was performed. Here, NPXc i.e. LMG8989, LMG8992 and LMG9002 were having approx. 93% and 50% ANI and dDDH values with *X. sontii* strains respectively. Hence, none of the NPXc are from species *X. sontii*. Moreover, these three strains amongst themselves were having ANI ranging from 92.7% to 93.7% and dDDH ranging from 48.9% to 52.7%. Hence, all these three strains represent three different novel genomospecies, which we are referring as NPXc complex. The NPXc complex is clubbing with *X. sontii* which is reported earlier as non-pathogenic species from rice (Bansal, Kaur et al. 2019). Interestingly, fourth NPXc isolate, LMG8993, and walnut associated NPXw strains (CFBP7634 and CFBP7651) were having around 96% orthoANI but 67% dDDH values. All these NPX strains were having around 96% orthoANI but dDDH ranged from 67% to 70%. Hence, according to multiple genome similarity parameters, these three NPX isolates form a potential taxonomic outlier of pathogenic *X. arboricola* (figure 2). While NPXc and NPXr isolates are in both clades I and II, the NPXw isolates are only in clade I.

**Figure 1:**
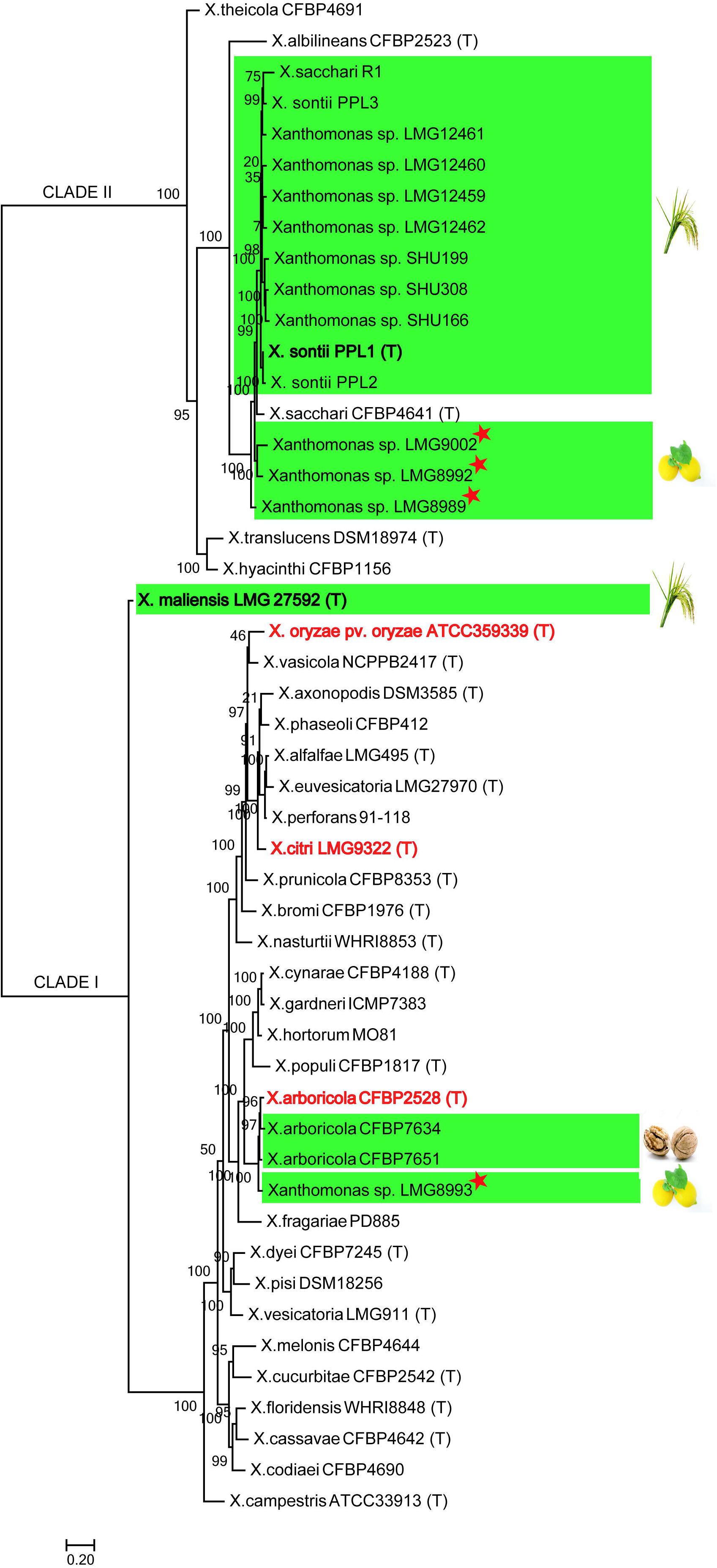
PhyloPhlAn tree of non-pathogenic and pathogenic *Xanthomonas* constructed using core genome. Here, all the NPX strains are highlighted in green box and their host are shown. Pathogenic species *X. citri, X. oryzae* and *X. arboricola* are highlighted in red. The type strains (designated as (T)) and strains sequenced in the study are represented by stars.

**Figure 2:**
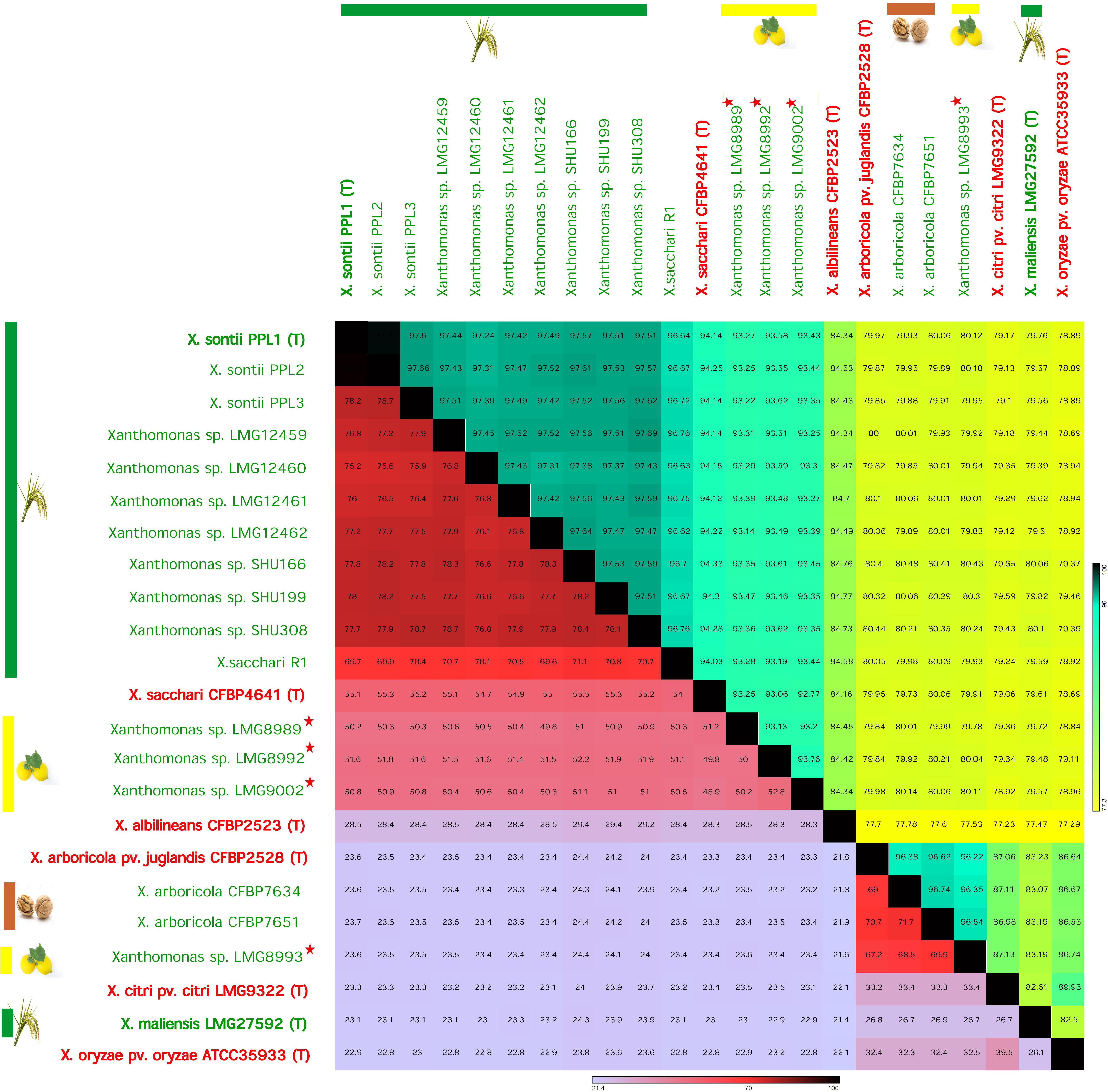
Taxonogenomics of all NPX. Non-pathogenic and pathogenic strains are in red and green color. Non-pathogenic strains from rice, lemon and walnut are represented by green, yellow and brown bars. The type strains are designated as (T) and strains sequenced in the study are represented by stars. Here, OAI values are represented in upper triangle and dDDH values are in lower triangle.

### Virulence determining clusters among different lifestyle bacteria

Genome-wide studies of these ecological and phylogenomic relatives with diverse lifestyles allow us to understand the evolution of highly successful lineages. Hence, we compared the status of well-characterized pathogenicity related gene(s) or gene cluster(s) in these non-pathogenic isolates/species from diverse hosts belonging to both the clades (figure 3). Overall the analysis revealed that the key regulators of pathogenicity were found to be absent in non-pathogenic isolates from clade II with a few exceptions from clade I. Global regulators of pathogenicity i.e. *hrpG* and *hrpX* were absent from all non-pathogenic strains of clade II and present in those from clade I. In contrast, rich repertoires of cell wall degrading enzymes and adhesins were observed for NPX strains of clade I as compared to clade II. Among adhesins, *pilQ, xadA* and *xadB* were present in all the NPX strains irrespective of their clades and *yapH* was present only in NPX isolates from clade I.

**Figure 3:**
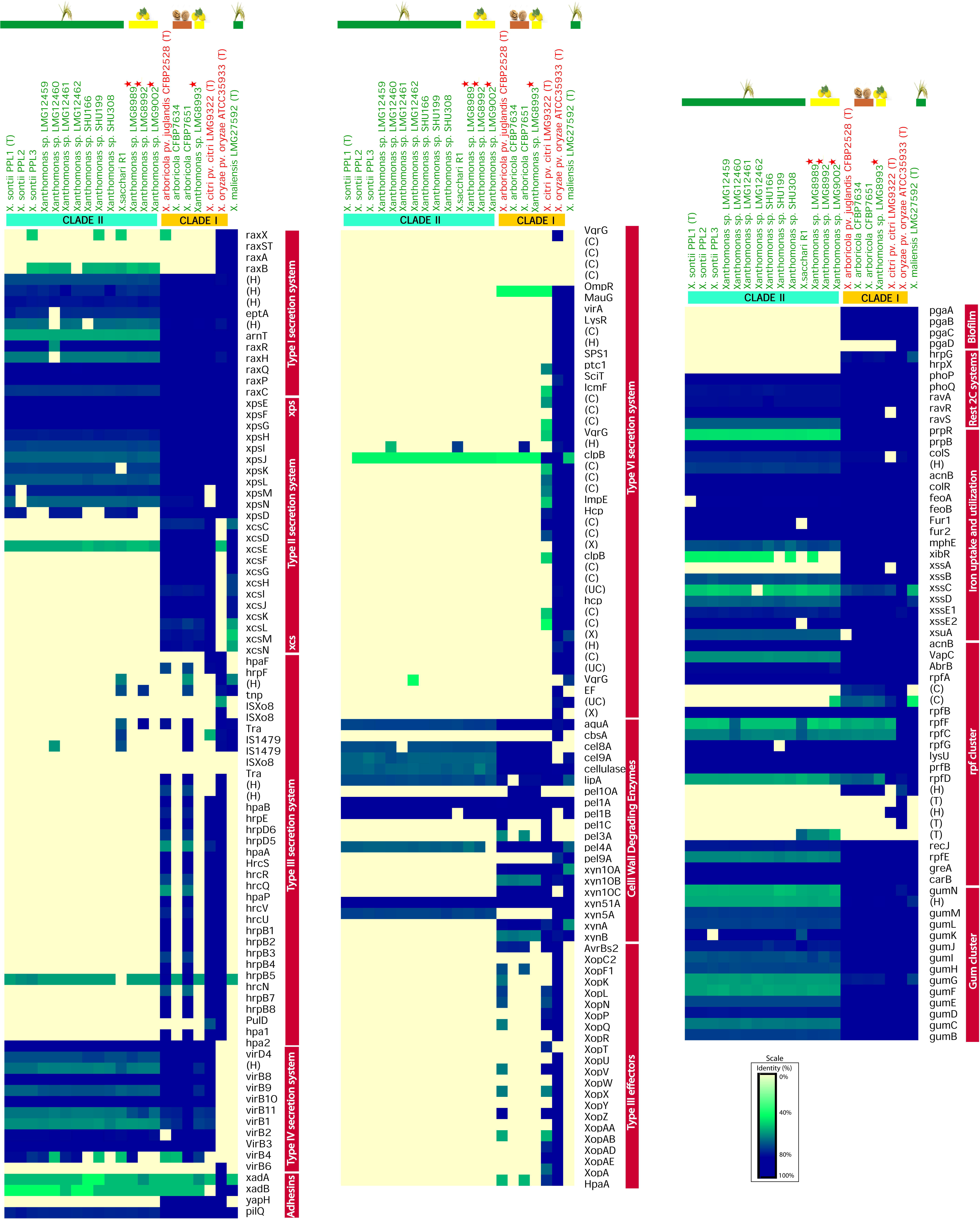
Heatmap showing status of various pathogenicity related clusters. (names indicated in vertical boxes) for non-pathogenic strains in the study and their respective pathogenic counterparts. Intensity of the color indicates level of identity with the query as depicted by the scale.

Amongst pathogenicity related gene clusters, T3SS, their effectors, T6SS and biofilm cluster were absent from all the non-pathogenic isolates from clade II irrespective of their host of isolation. However, there are few exceptions in NPX isolates from clade I. One of the NPXc (LMG8993) and NPXw isolates had biofilm cluster (except for *pgaD*). One of the NPXw (CFBP7651) have a non-canonical T3SS in accordance to a comparative genomic study pathogenic and non-pathogenic *X. arboricola* isolates (Essakhi, Cesbron et al. 2015). Furthermore, *X. maliensis*, eventhough lacking T3SS, was having key pathogenicity related gene clusters like T6SS and biofilm forming clusters.

Among other secretion systems, T1SS is present in all the NPX isolates, except for *raxST* and *raxA* absent from all the isolates except for *X. oryzae* and *X. maliensis*. Further, *raxX* and *raxB* were also found to be diversifying or absent in all except *X. oryzae* and *X. maliensis*. Among the T2SS, *xps* is present in all the isolates however, *xcs* is specific to clade I NPX isolates. The T4SS was present in all the NPX isolates except for *X. maliensis*. Further, commonality in the gene clusters like: iron uptake and utilization related genes, *rpf* cluster, *gum* cluster and *xps* cluster etc. in all the strains irrespective of their lifestyles.

## Discussion

Pathogenic strains that infect citrus plants belong to a single species (Bansal, Midha et al. 2017). In this context, it is interesting to find that non-pathogenic isolates from *Citrus* sp. are highly diverse with each isolate belonging to a novel genomospecies and altogether forming a species complex, NPXc complex. This suggests existence of a highly diverse and co-evolving community of NPX on citrus plants. This is unlike NPX isolates from rice that majorly belong to one species *X. sontii*. Owing to the diversity of NPXc, there is a need to expand such study by isolating and investigating large number of NPX strains from citrus plants. Incidentally, unlike rice, citrus plants themselves belong to several highly diverse species. Hence, such studies are required to be expanded to all citrus species and not just one species to understand origin, evolution and adaptation of NPX population on citrus hosts.

Phylogenetically pathogenic species of rice (*X. oryzae*) and citrus (*X. citri* pv. citri) are distantly related (Midha and Patil 2014) and unlike NPX isolates, both of the pathogenic species fall in a single clade (clade I). This may be to do with the pathogenic lifestyle on respective hosts and requirement of distinct set of virulence genes. Eventhough, the NPX isolates are from two diverse hosts (monocot vs dicot) but in phylogenomic tree majority of the isolates club together as closely related species group. This highlights origin of both the communities from a common ancestor in recent past and that lifestyle is also important along with host specialization. There is parallel evolution and convergence of function related to this lifestyle. This also means requirement of core set of functions to be a successful non-pathogenic *Xanthomonas*.

Most of the NPX isolates in both rice and citrus belong to clade II. At the same time minority of the strains in both rice (*X. maliensis*), citrus (LMG8993) and even *Xanthomonas* associated with walnut (NPXw, CFBP7634 and CFBP7651) were found to be more related to pathogenic isolates. This suggests existence of minor group that may be ancestral to the pathogenic counterparts and ongoing dynamics towards diverse lifestyles. Also, pathogenic *X. citri* and *X. arboricola* species are known to cause similar type of symptoms in diverse hosts (Garita-Cambronero, Sena-Vélez et al. 2019). Clubbing of a NPX isolate from citrus with pathogenic and non-pathogenic isolates of walnut indicates basic functions required to adapt to fruit plants. Moreover, previous study on comparison of pathogenic and non-pathogenic strains from walnut found to belong to the same species (*X. arboricola*). Interestingly, whole genome based comparison have shown that NPXw isolates (CFBP7651, also included in the present study) were having non-canonical T3SS. Similarly, amongst NPXr as well *X. maliensis* was found to be more related to pathogenic isolates and also having some of the key pathogenicity determinants which are otherwise absent from majority of NPX isolates (Bansal, Midha et al. 2019).

Hence, present study reflects that phylogenomically and gene-content wise, NPX have two waves of evolution. First one is followed by majority of NPX and forming an NPX complex with absence of virulence related clusters. While, the second one are more diversified and related to pathogenic strains, although, they are represented by minority of the strains. Knowledge pertaining to the population of NPX from diverse hosts will shed light on epidemiology and evolution of pathogenic isolates. Genomics has opened a new era in *Xanthomonas* research and large scale on both minor and major groups of NPX isolates across diverse hosts likes grasses, shrubs, trees, etc. is warranted.

## Material and methods

### Strain collection, genome sequencing, assembly and annotation

All the four *Citrus* sp. associated strains were collected from BCCM-LMG culture collection and were grown as per the instructions. Genomic DNA was extracted by using a ZR Fungal/Bacterial DNA MiniPrep kit (Zymo Research). DNA quality checking was done using a NanoDrop 1000 instrument (Thermo Fisher Scientific) and agarose gel electrophoresis. Quantitation of DNA was performed using a Qubit 2.0 fluorometer (Life Technologies). Illumina paired-end sequencing libraries (read length, 2 × 250) of genomic DNA were prepared using Nextera XT sample preparation kits (Illumina, Inc., San Diego, CA, USA) with dual indexing adapters. Illumina sequencing libraries were sequenced in-house using an Illumina Miseq platform (Illumina, Inc., San Diego, CA, USA) and company-supplied paired-end sequencing kits. Adapter trimming was done automatically by MiSeq control software (MCS), and additional adapter contamination identified by the NCBI server was removed by manual trimming. Raw reads were assembled *de novo* using CLC Genomics Workbench v7.5 (CLC bio, Aarhus, Denmark) with default settings. Annotation was done using the NCBI PGAP pipeline.

### Phylogenomics and taxonogenomics

PhyloPhlAn v0.99 was used to construct the phylogenomic tree based on more than 400 conserved genes from whole genome proteome data (Segata, Börnigen et al. 2013). USEARCH v5.2.32 (Edgar 2010), MUSCLE v3.8.31 (Edgar 2004) and FastTree v2.1 (Price, Dehal et al. 2009) were used for ortholog searching, multiple sequence alignment and phylogenomic construction respectively. Taxonogenomic analysis of the strains was performed using OAI calculated using USEARCH v5.2.32 and dDDH values were calculated using genome to genome distance calculator (http://ggdc.dsmz.de/distcalc2.php).

### Virulence related gene(s) and gene cluster(s) analysis

Gene clusters related to virulence were extracted (Bansal, Midha et al. 2019) and tBLASTn searches were performed using standalone BLAST+ 2.9.0. Cut-offs used for similarity and coverage were 40% and 50% respectively. Heatmaps for the blast searches were generated using GENE-E v3.03215 (https://software.broadinstitute.org/GENE-E/).

## Author Contributions

KB and SK have performed genome sequencing. SK performed phylogenomic and comparative analysis. KB have performed taxonogenomic and virulence gene cluster analysis. KB have drafted manuscript with inputs from SK, and PBP. PBP conceived the study and participated in its design with KB. All the authors have read the manuscripts and approved the manuscript.

## Acknowledgements

This work is supported by a project entitled “Megagenomics and Metagenomics insights into adaptation and evolution of fruit Microbiome” (GAP0187) and “(MICRA) – Mega-genomic insights into co-evolution of rice and its Microbiome” (MLP0020) to PBP.

---

**Supplementary table 1:**
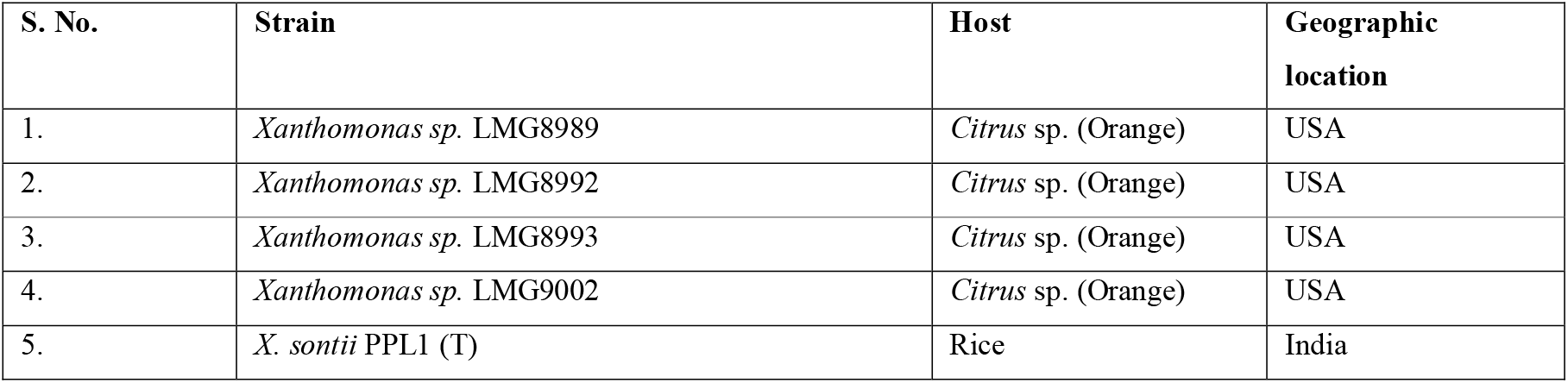
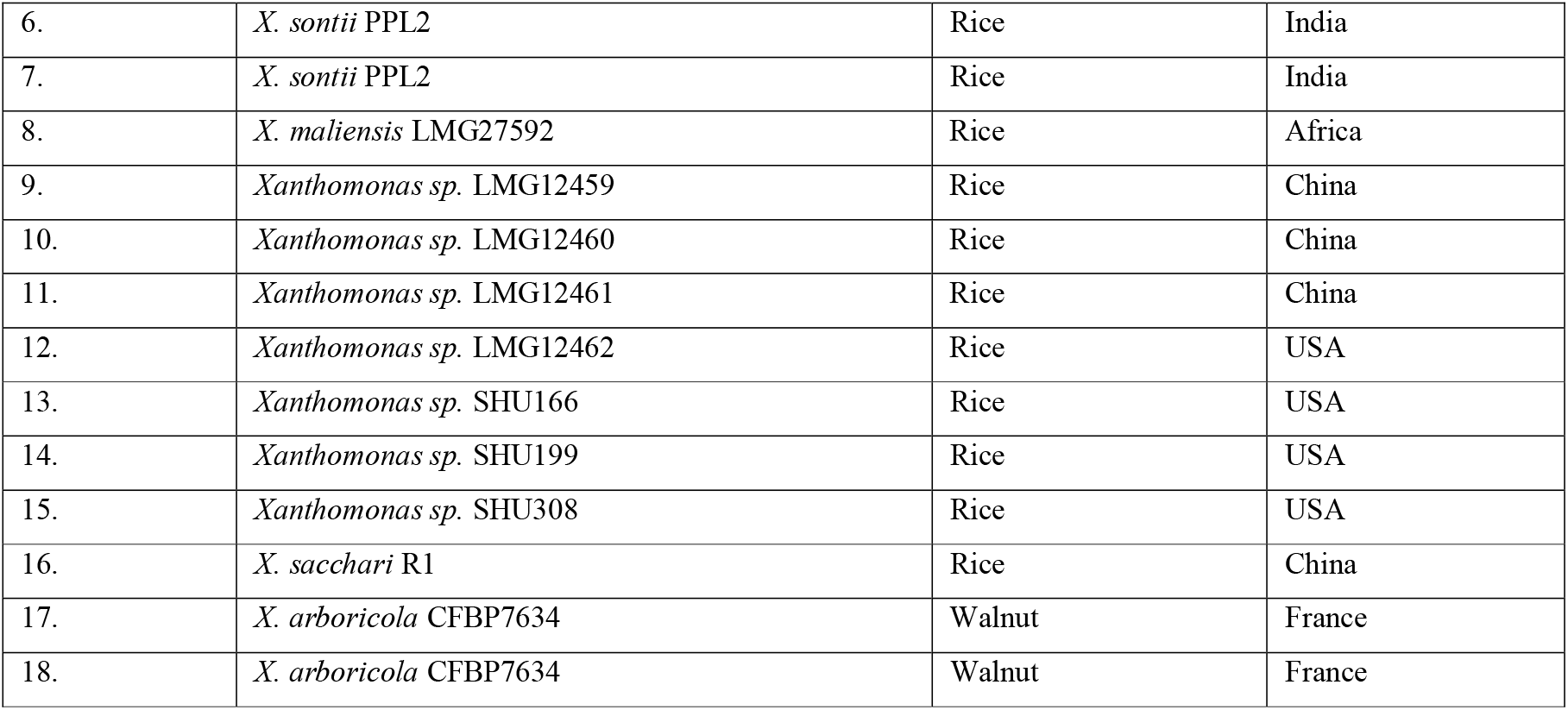
Metadata of non-pathogenic strains used in the study.

## References

Bansal, K., et al. (2019). “Xanthomonas sontii sp. nov., a non-pathogenic bacterium isolated from healthy basmati rice (Oryza sativa) seeds from India.” bioRxiv: 738047.

Bansal, K., et al. (2019). “Ecological and evolutionary insights into pathogenic and non-pathogenic rice associated Xanthomonas.” bioRxiv: 453373.

Bansal, K., et al. (2017). “Ecological and evolutionary insights into Xanthomonas citri pathovar diversity.” Applied and environmental microbiology 83(9): e02993–02916.

Bernal, P., et al. (2018). “Type VI secretion systems in plant□associated bacteria.” Environmental microbiology 20(1): 1–15.

Brunings, A. M. and D. W. Gabriel (2003). “Xanthomonas citri: breaking the surface.” Molecular Plant Pathology 4(3): 141–157.

Büttner, D. and U. Bonas (2010). “Regulation and secretion of Xanthomonas virulence factors.” FEMS microbiology reviews 34(2): 107–133.

Cesbron, S., et al. (2015). “Comparative genomics of pathogenic and nonpathogenic strains of Xanthomonas arboricola unveil molecular and evolutionary events linked to pathoadaptation.” Frontiers in plant science 6: 1126.

Chan, J. W. and P. H. Goodwin (1999). “The molecular genetics of virulence of Xanthomonas campestris.” Biotechnology advances 17(6): 489–508.

Chatterjee, S. and R. V. Sonti (2002). “rpfF mutants of Xanthomonas oryzae pv. oryzae are deficient for virulence and growth under low iron conditions.” Molecular Plant-Microbe Interactions 15(5): 463–471.

Dow, J. M., et al. (2000). “Novel genes involved in the regulation of pathogenicity factor production within the rpf gene cluster of Xanthomonas campestris.” Microbiology 146(4): 885–891.

Edgar, R. C. (2004). “MUSCLE: multiple sequence alignment with high accuracy and high throughput.” Nucleic acids research 32(5): 1792–1797.

Edgar, R. C. (2010). “Search and clustering orders of magnitude faster than BLAST.” Bioinformatics 26(19): 2460–2461.

Essakhi, S., et al. (2015). “Phylogenetic and variable-number tandem-repeat analyses identify nonpathogenic Xanthomonas arboricola lineages lacking the canonical type III secretion system.” Appl. Environ. Microbiol. 81(16): 5395–5410.

Etchegaray, A., et al. (2004). “In silico analysis of nonribosomal peptide synthetases of< i> Xanthomonas axonopodis</i> pv.< i> citri</i>: identification of putative siderophore and lipopeptide biosynthetic genes.” Microbiological research 159(4): 425–437.

Garita-Cambronero, J., et al. (2019). “Xanthomonas citri subsp. citri and Xanthomonas arboricola pv. pruni: Comparative analysis of two pathogens producing similar symptoms in different host plants.” PLoS One 14(7): e0219797.

Gottwald, T. R., et al. (2001). “The citrus canker epidemic in Florida: the scientific basis of regulatory eradication policy for an invasive species.” Phytopathology 91(1): 30–34.

Gottwald, T. R., et al. (2002). “Geo-referenced spatiotemporal analysis of the urban citrus canker epidemic in Florida.” Phytopathology 92(4): 361–377.

Graham, J. H., et al. (2004). “Xanthomonas axonopodis pv. citri: factors affecting successful eradication of citrus canker.” Molecular Plant Pathology 5(1): 1–15.

Hayward, A. (1993). The hosts of Xanthomonas. Xanthomonas, Springer: 1–119.

Jha, G., et al. (2007). “Functional interplay between two Xanthomonas oryzae pv. oryzae secretion systems in modulating virulence on rice.” Molecular plant-microbe interactions 20(1): 31–40.

Kumar, S., et al. (2019). “Phylogenomics insights into order and families of Lysobacterales.” Access Microbiology 1(2).

Lee, S.-W., et al. (2008). “The Xanthomonas oryzae pv. oryzae PhoPQ two-component system is required for AvrXA21 activity, hrpG expression, and virulence.” Journal of Bacteriology 190(6): 2183–2197.

Leyns, F., et al. (1984). “The host range of the genusXanthomonas.” The Botanical Review 50(3): 308–356.

Lu, H., et al. (2008). “Acquisition and evolution of plant pathogenesis-associated gene clusters and candidate determinants of tissue-specificity in xanthomonas.” PLoS One 3(11): e3828.

Midha, S., et al. (2017). “Population genomic insights into variation and evolution of Xanthomonas oryzae pv. oryzae.” Scientific reports 7: 40694.

Midha, S. and P. B. Patil (2014). Genomic Flux in Xanthomonas Group of Plant Pathogenic Bacteria. Plasticity in Plant-Growth-Promoting and Phytopathogenic Bacteria. E. I. Katsy. New York, Springer: 131–153.

Pandey, A. and R. V. Sonti (2010). “Role of the FeoB protein and siderophore in promoting virulence of Xanthomonas oryzae pv. oryzae on rice.” Journal of Bacteriology 192(12): 3187–3203.

Price, M. N., et al. (2009). “FastTree: computing large minimum evolution trees with profiles instead of a distance matrix.” Molecular biology and evolution 26(7): 1641–1650.

Qian, W., et al. (2008). “Genome-scale mutagenesis and phenotypic characterization of two-component signal transduction systems in Xanthomonas campestris pv. campestris ATCC 33913.” Mol Plant Microbe Interact 21(8): 1128–1138.

Segata, N., et al. (2013). “PhyloPhlAn is a new method for improved phylogenetic and taxonomic placement of microbes.” Nature communications 4: 2304.

Subramoni, S., et al. (2012). “The ColRS system of Xanthomonas oryzae pv. oryzae is required for virulence and growth in iron□limiting conditions.” Molecular Plant Pathology 13(7): 690–703.

Triplett, L. R., et al. (2015). “Characterization of a novel clade of Xanthomonas isolated from rice leaves in Mali and proposal of Xanthomonas maliensis sp. nov.” Antonie Van Leeuwenhoek 107(4): 869–881.

Tsuge, S., et al. (2006). “Gene involved in transcriptional activation of the hrp regulatory gene hrpG in Xanthomonas oryzae pv. oryzae.” Journal of Bacteriology 188(11): 4158–4162.

Vauterin, L., et al. (1996). “Identification of non-pathogenic Xanthomonas strains associated with plants.” Systematic and Applied Microbiology 19(1): 96–105.

Wengelnik, K. and U. Bonas (1996). “HrpXv, an AraC-type regulator, activates expression of five of the six loci in the hrp cluster of Xanthomonas campestris pv. vesicatoria.” Journal of Bacteriology 178(12): 3462–3469.

Wengelnik, K., et al. (1999). “Mutations in the Regulatory Gene hrpG ofXanthomonas campestris pv. vesicatoria Result in Constitutive Expression of All hrp Genes.” Journal of Bacteriology 181(21): 6828–6831.

