## supplementary table 1 for "Extreme phylogenomic diversity in non-pathogenic *Xanthomonas* strains associated with citrus"

**Supplementary table 1:** Metadata of non-pathogenic strains used in the study.

| **S. No.** | **Strain** | **Host** | **Geographic location** |
| --- | --- | --- | --- |
| 1. | *Xanthomonas sp.* LMG8989 | *Citrus* sp. (Orange) | USA |
| 2. | *Xanthomonas sp.* LMG8992 | *Citrus* sp. (Orange) | USA |
| 3. | *Xanthomonas sp.* LMG8993 | *Citrus* sp. (Orange) | USA |
| 4. | *Xanthomonas sp.* LMG9002 | *Citrus* sp. (Orange) | USA |
| 5. | *X. sontii* PPL1 (T) | Rice | India |
| 6. | *X. sontii* PPL2 | Rice | India |
| 7. | *X. sontii* PPL2 | Rice | India |
| 8. | *X. maliensis* LMG27592 | Rice | Africa |
| 9. | *Xanthomonas sp.* LMG12459 | Rice | China |
| 10. | *Xanthomonas sp.* LMG12460 | Rice | China |
| 11. | *Xanthomonas sp.* LMG12461 | Rice | China |
| 12. | *Xanthomonas sp.* LMG12462 | Rice | USA |
| 13. | *Xanthomonas sp.* SHU166 | Rice | USA |
| 14. | *Xanthomonas sp.* SHU199 | Rice | USA |
| 15. | *Xanthomonas sp.* SHU308 | Rice | USA |
| 16. | *X. sacchari* R1 | Rice | China |
| 17. | *X. arboricola* CFBP7634 | Walnut | France |
| 18. | *X. arboricola* CFBP7634 | Walnut | France |
